# Cellular Migration May Exhibit Intrinsic Left-Right Asymmetries: A meta-analysis

**DOI:** 10.1101/269217

**Authors:** Kelly G. Sullivan, Laura N. Vandenberg, Michael Levin

**Affiliations:** Biology Department, Center for Regenerative and Developmental Biology, Tufts University, Medford, MA 02155; Department of Public Health, Division of Environmental Health Sciences, University of Massachusetts – Amherst, Amherst, MA 01003

**Keywords:** cilia, intracellular, ion transporter, laterality, microtubule, chirality, asymmetry, motility

## Abstract

The intracellular model of embryonic left-right (LR) asymmetry proposes that body laterality originates from intrinsic chiral properties of individual cells, and several recent studies identified consistent chirality in the behavior of cells in vitro. Here, we explored one prediction of the intrinsic asymmetry model: that LR asymmetries would be present in a wide range of mammalian cells, manifesting in the form of LR-biased migration toward an attractant. We mined data from published papers on galvanotaxis and chemotaxis and quantitatively analyzed the migration trajectories of adult somatic cells, stem cells, and cancer cells to determine whether they display significant consistent LR biases in their movements toward migration targets. We found that several cell types exhibited LR biases during galvanotaxis and chemotaxis, and that treatments inhibiting cytoskeletal remodeling or targeting ion channel activity both abolished these LR biases. While we cannot conclusively rule out the existence of subtle biasing cues in the apparatus of some of the studies, the analysis of this dataset suggests specific assays and cell types for further investigation into the chiral aspects of intrinsic cell behavior.

**Funding:** This work was funded by the American Heart Association Established Investigator grant 0740088N and NIH grants R01-GM077425 (to ML) and NRSA grant 1F32GM087107 (to LNV). M.L. is also supported by the G. Harold and Leila Y. Mathers Charitable Foundation.

## Introduction

Although most vertebrates have external bodies that appear symmetric along the left-right (LR) axis, these same animals have asymmetric internal body plans. These biases encompass not only the placement and morphology of single organs such as the heart, spleen and stomach, but also organs such as the lungs, which have distinct morphologies on the left and right sides (Raya and Belmonte, 2006; Vandenberg and Levin, 2010). Establishing LR asymmetry is an important aspect of normal embryonic development, and diseases of laterality affect approximately 1 in 8,000 births (Casey and Hackett, 2000; Cohen et al., 2007; Hackett, 2002; Peeters and Devriendt, 2006). Understanding how and when the embryo reliably distinguishes its left from its right is an important question for developmental and cell biology, evolution, and the biomedicine of birth defects.

Three major models of LR asymmetry have been proposed, each with distinguishing characteristics that can be assessed experimentally (Levin and Palmer, 2007; Speder et al., 2007; Vandenberg et al., 2013a; Vandenberg and Levin, 2013). The first model, termed the ciliary model, proposes that the LR axis is established in the late gastrula and early neurula stages of development due to the movement of cilia that produce a LR-biased fluid flow within a small fluid-filled space in the embryo, often called the node (Basu and Brueckner, 2008; Komatsu and Mishina, 2013; Tabin, 2005; Yost, 2003). The second model, termed the chromatid segregation model, postulates the consistently asymmetric distribution of differentially imprinted chromatids during the first cleavage stage (Armakolas and Klar, 2007; Sauer and Klar, 2012), establishing the molecular distinction of the left and right sides from the earliest stages of development (Klar, 1994; Klar, 2008; Sauer and Klar, 2012). The third model, termed the intracellular model, also proposes that the LR axis is established very early in development (Aw and Levin, 2008; Levin and Nascone, 1997; Levin and Palmer, 2007). In this model, a chiral cytoskeleton orients the LR axis with respect to the anterior-posterior and dorso-ventral axes, and actively directs the asymmetric intracellular localization of proteins, including K^+^ channels and H^+^ pumps, in the early cleavage stage embryo (Aw et al., 2008; Lobikin et al., 2012; McDowell et al., 2016a; McDowell et al., 2016b; Qiu et al., 2005; Vandenberg et al., 2013a; Vandenberg and Levin, 2013; Vandenberg et al., 2013b). These biases in ion transporter localization produce consistent biases in the pH and transmembrane voltage on the left and right sides of the embryo (Adams et al., 2006; Morokuma et al., 2008), which drive the asymmetric distribution of charged signaling molecules, such as pre-nervous serotonin, across large cell fields (Carneiro et al., 2011; Fukumoto et al., 2003; Fukumoto et al., 2005). The intracellular model leverages asymmetric signaling across large cell fields from the intracellular chirality of early blastomeres, acting as a biophysical amplifier of single cell asymmetries.

The intracellular models predict that consistent chirality is an ancient, universal property of single cells (Davison et al., 2016; Naganathan et al., 2016; Raymond et al., 2017; Tamada and Igarashi, 2017; Tee et al., 2015). While this is known to be the case for unicellular ciliates (Frankel, 1991) and slime molds (Dimonte et al., 2016), and increasingly being discovered in snail, nematode, and fruit fly embryos (Callander et al., 2014; Chen et al., 2016; Davison et al., 2016; Hayashi and Murakami, 2001; Inaki et al., 2016; Kuroda et al., 2009; Naganathan et al., 2014; Sato et al., 2015; Schonegg et al., 2014; Taniguchi et al., 2011), it is still often claimed that asymmetry in vertebrates requires a large, ciliated node structure that can support strong fluid flows. Interestingly, several studies have reported that individual cells can exhibit consistent chiral behaviors in culture in the direction of neurite outgrowth, cell:substrate interactions, and docking with neighboring cells (Chen et al., 2012; Frankel, 1991; Heacock and Agranoff, 1977; Nelsen et al., 1989; Wan et al., 2011; Wan and Vunjak-Novakovic, 2011). Wildtype HL-60 neutrophil-like cells, for example, typically extend a pseudopod to the left of the nucleus-centrosome axis following exposure to fMLP, a chemoattractant (Xu et al., 2007). This leftward bias can be ablated with chemicals that disrupt the cytoskeleton or molecular genetic disruptions to α-tubulin (Lobikin et al., 2012; Xu et al., 2007).

We performed a meta-analysis testing two hypotheses: 1) that mammalian cells exhibit LR biased migration paths, and 2) that these asymmetries would be observable in a wide range of cell types. Migration plays crucial roles in embryonic development, and thus asymmetries in migration behavior of single cells are expected to impact large-scale morphogenesis (Lenhart et al., 2013; Rohr et al., 2008; Yost, 1990). We examined published studies of galvanotaxis and chemotaxis and assessed cell migration trajectories to determine whether cultured cells experience consistent LR deviations in cell movement towards an attractant. Whereas the null hypothesis predicts an equal number of cells deviating to the right vs. left of a straight line of migration, we observed consistently LR-biased migration in a range of different cell types. We also analyzed published results of migration experiments in cells treated with reagents that inhibit cytoskeletal remodeling or alter ion flux and/or membrane potential. Our analysis reveals that LR biases in several mammalian cell types can be disrupted with treatments similar to those that abolish LR biases in embryos, reinforcing the importance of intrinsic chirality processes at the cellular level.

## Results & Discussion

### Apparent LR biases in cell migration occur in response to chemical and electrical stimuli

A number of cell types have migratory responses to chemical and electrical gradients, both in culture and in the developing embryo (Chuai et al., 2012; Rorth, 2002; Thiery, 1984; Zhao, 2009), but to our knowledge no one has examined whether they exhibit LR-biased movements. To assess whether these migratory responses have inherent LR biases, we analyzed studies examining galvanotactic or chemotactic responses of cultured cells (Fig. 1). A number of cells were shown to have left-or right-biased migration patterns following treatment with stimuli (Table 1). Interestingly, many of these LR-biases were apparent only under a narrow range of conditions (i.e. stimulation of cells isolated from the bovine outer meniscus treated with 2, 4 or 6 V/cm, but not 0.2 or 0.8 V/cm) (Gunja et al., 2012), suggesting that the extent of the bias we found may be an underestimate: cell types which revealed no bias in a particular study may exhibit biases when challenged by different conditions.

**Figure 1.**
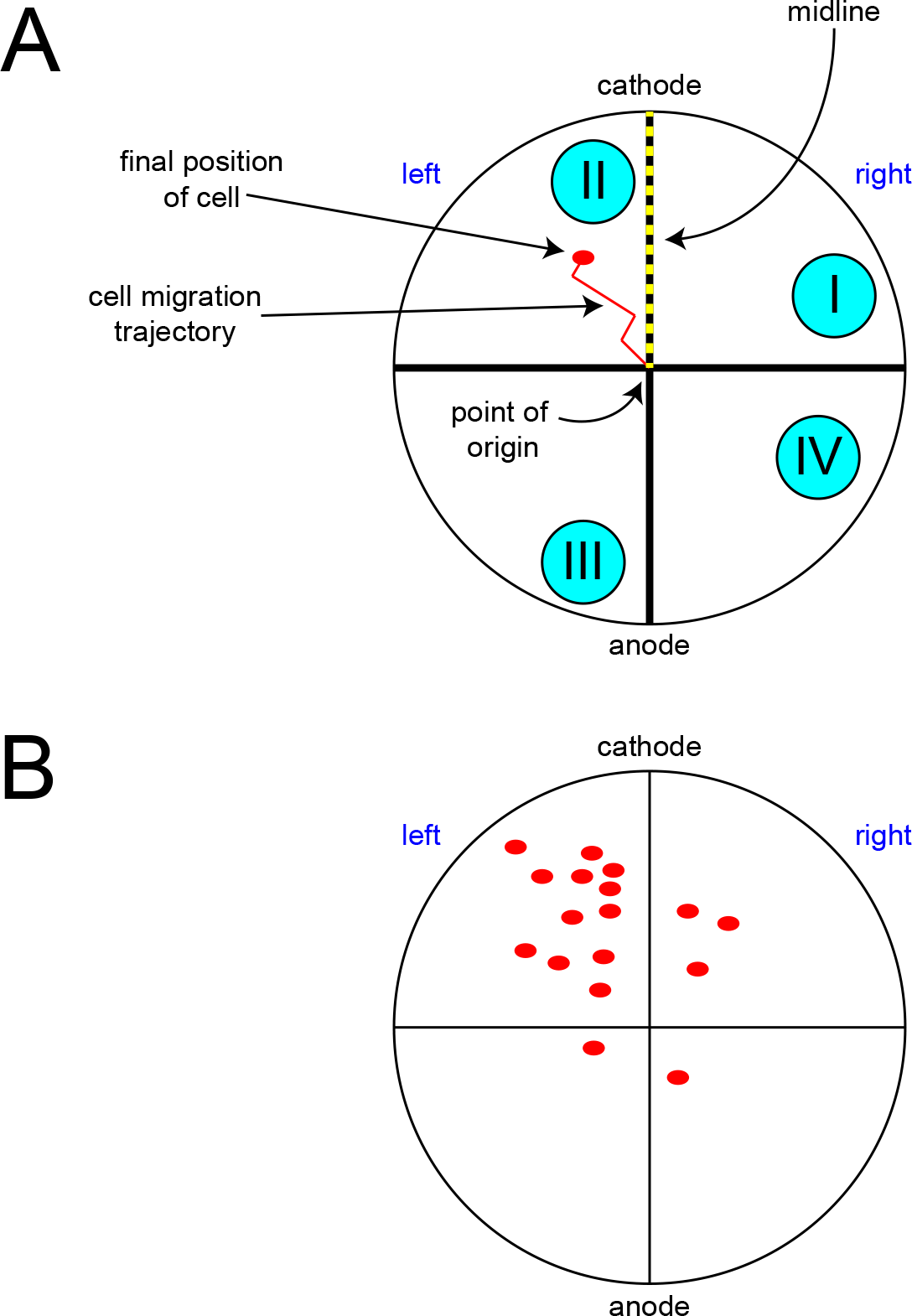
Assessment of LR biases in cell migration. A) A sample Cartesian plot. The original locations of all cells were placed on a common origin at the center of the plot in the cited studies. The galvanotactic or chemotactic treatment was applied to attract cells toward the “North” direction on the culture plate (indicated here by the cathode as an example). The axis of cell attraction (dotted line indicating the midline) and the axis perpendicular to the dish surface (the dish vs. media or “dorso-ventral” axis) establish two orthogonal axes with respect to which a third LR axis can be defined. The trajectories and/or final positions of cells are indicated in each study and scored by our meta-analysis to determine whether they ended up to the left or right of the midline (null hypothesis predicting no net bias in each experiment overall). Only cells that migrated to quadrants I and II were counted for analyses; cells that migrated to quadrants III and IV are considered non-responsive (or insensitive) to the applied treatment. B) A sample plot illustrating data for cells that exhibit a leftward bias with respect to the migration line.

**Table 1:** LR biases are observed in the migratory patterns of cultured cells

| Study | Cell type | G/C<sup>a</sup> | Treatments inducing LR bias | Direction of bias | p-value |
| --- | --- | --- | --- | --- | --- |
| (Pullar et al., 2001). | Human keratinocytes | G | 100mV/mm | Right | < 0.01 |
| (Nishimura et al., 1996). | Human Keratinocytes | G | 100mV/mm, 200 mV/mm | Right | < 0.05 |
| (Gunja et al., 2012) | bovine outer knee meniscus | G | 2V/cm, 4V/cm, 6V/cm | Left | < 0.001 |
|  | bovine inner knee meniscus | G | 0.2V/cm, 2V/cm, 4V/cm, 6V/cm | Left | < 0.01 |
| (Messerli and Graham, 2011) | zebrafish keratocytes | G | 10mV/mm, 100mV/mm | Left | < 0.001 |
| (Li et al., 2011) | human T-cells | C | CCL19 | Right | < 0.001 |
| (Sackmann et al., 2012) | Human primary neutrophils (Type 2) | C | FMLP | Right | < 0.001 |
| (Zicha et al., 1998) | WAS neutrophils | C | FMLP | Left | < 0.001 |
|  | WAS macrophages | C | CSF1 | Left | < 0.001 |
| (Li et al., 2012) | MDCK II (monolayer) | G | 200mV/mm | Right | < 0.001 |
|  | NRK (isolated) | G | 200 mV/mm | Left | <0.01 |
|  | NRK (monolayer) | G | 200mV/mm | Right | < 0.001 |
|  | Tracheal epithelial cells (monolayer) | G | 200mV/mm | Right | < 0.001 |
| (Barbet et al., 2008) | iDC (dendritic) | C | CCL21 | Right | < 0.001 |
| (Feng et al., 2012) | human neural stem cells (from H9 line) | G | 16mV/mm, 250mV/mm, 300mV/mm | Left | < 0.01 |
| (Meng et al., 2011) | rat aNPC | G | 500 mV/mm | Left | < 0.001 |
| (Rovasio et al., 2012) | chick neural crest cells | C | SCF | Left | < 0.01 |
| (Yao et al., 2008) | rat hippocampal neurons | G | 200 mV/mm, 300mV/mm | Left | < 0.05 |
| (Zhao et al., 2011) | Human BM-MSC | G | 200mV/mm | Left | < 0.001 |
| (Djamgoz et al., 2001) | Mat-LyLu | G | 3V/cm | Left | < 0.001 |
|  | AT-2 |  | 3V/cm | Right | < 0.001 |
| (Martin-Granados et al., 2012) | HeLa-Tet-Off | G | 200mV/mm | Right | < 0.001 |
|  | PC3-3-M | G | 200mV/mm | Left | < 0.001 |
| (Pu et al., 2007) | MTLn3 | G | 1V/cm | Left | < 0.01 |
| (Raja et al., 2010) | MTLn3 Mena<sup>Inv</sup> | C | 5.5 μM EGF | Left | < 0.05 |
| (Sun et al., 2012) | CL1-5 | G | 338mV/mm | Left | < 0.05 |
| (Wang et al., 2002) | PLB-985 | C | C5a | Right | < 0.001 |
| (Wu et al., 2013) | MDA-MB-231 | G | 1.5V/cm | Left | <0.001 |
| (Hwang et al., 2014) | SKOV3 | C | LPA | Right | < 0.05 |
| (Gao et al., 2011) | Dictyostelium | G | 12V/cm pH 6.5 | Right | < 0.05 |
| (Sato et al., 2009) | Dictyostelium | G | 10V/mm | Left | < 0.001 |

<sup>a</sup> G = galvanotaxis, C = chemotaxis. Highlighted rows are cancer cells. Additional information about treatments that did not induce LR biases is presented in Supplemental Table 1.

Of the distinct cell types displaying LR biases after galvanotactic or chemotactic treatments, approximately half were biased on the left and half to the right (Table 1). Thus, while possession of consistent asymmetry is a common feature of many cell types, the direction of their preferred asymmetry is not universal. Left-biased cells included those isolated from connective tissue, some neural cells and stem cells. Right-biased cells included keratinocytes, epithelial cells, some neural cells, and immune cells. Unbiased cells were largely represented by fibroblasts and related cell types (Supplemental Table 1). Several cancer cell lines examined in galvanotaxis or chemotaxis assays also displayed a LR bias, including cells isolated from lung, breast, cervical and prostate cancers and myeloid leukemia (Table 1). The fact that different cell types have opposite chirality is interesting, but not surprising given similar range of behavioral preferences of different cell types for stimuli such as example anodes vs. cathodes (Kim et al., 2017; Korohoda et al., 2000; Mycielska and Djamgoz, 2004; Sun et al., 2013; Van Hoek et al., 1999).

Collectively, these results suggest that individual cell types have inherent LR biases, consistent with the hypothesis that LR asymmetry is an ancient feature of cells and not dependent on fluid flow in specialized vertebrate embryonic organs of asymmetry (Levin and Palmer, 2007; Vandenberg and Levin, 2009; Vandenberg and Levin, 2010). One factor that prevented us from analyzing other studies of galvanotaxis and chemotaxis was the manner in which the results are reported. We expect that additional cells are likely to also display LR biases, at least under certain conditions, but those data must be reported based on relative movements from a common origin for these asymmetries to be revealed.

An important limitation of this analysis is that we had no way of identifying possible technical biases in the analyzed studies, for example subtle misalignment of the main axis of a galvanotaxis experiment’s data plot with the actual field lines. In the absence of being able to rule out such possibilities, our meta-analysis is not conclusive with respect to any one cell type. However, the abolition of measured chirality by various chemical or genetic perturbations suggests that it is unlikely that experimental biases explain the entire effect at the meta-study level. We thus suggest our analysis as not a definitive proof of this new aspect of intrinsic chirality but rather as a providing a useful roadmap identifying the most promising specific cell types in specific assays for future investigations.

### LR biases are abolished with treatments that alter cytoskeletal remodeling, ion flux

We have previously proposed that the cytoskeleton could act as the origin of asymmetry in different cell types and different species, allowing single cells, plants, invertebrate animals, and vertebrates to use evolutionarily conserved mechanisms to establish a LR axis (Davison et al., 2016; Lobikin et al., 2012; McDowell et al., 2016a; McDowell et al., 2016b; Vandenberg et al., 2013a; Vandenberg and Levin, 2013). Prior studies suggest that the microtubule organizing center (MTOC) may offer an evolutionarily conserved structure by which asymmetry could be established in a wide range of species (Beisson and Jerka-Dziadosz, 1999; Bornens, 2012; Marshall, 2012). Because the MTOC is itself asymmetric, it could provide a means to amplify subcellular asymmetries to cell-and organ-level asymmetries (Mizuno et al., 2012), as it is known to do for planar cell polarity in general (Chen et al., 2016).

The studies we examined showed that specific cell types exhibited LR biases in their galvanotactic or chemotactic migratory responses. Our next analyses revealed that native biases could be abolished by pharmacological agents targeting specific pathways (Table 2). Consistent with our hypothesis that the cytoskeleton plays a central role in the establishment of LR asymmetry, drugs that targeted myosin or aspects of the Rac1/Rho signaling pathways, which are involved in the cytoskeletal remodeling necessary for cell migration, abolished LR biases (Feng et al., 2012; Pullar et al., 2001; Sato et al., 2009; Yao et al., 2008). Importantly, myosin has been implicated in LR patterning of vertebrate and invertebrate embryos (Nakamura et al., 2013; Noel et al., 2013), consistent with a role in asymmetries of both single cell migration and embryonic pattern formation (Hozumi et al., 2006; Petzoldt et al., 2012; Speder et al., 2006; Speder and Noselli, 2007).

**Table 2.** Abrogation of LR biases by additional treatments

| **Study** | **Cell type** | **Treatments inducing LR bias** | **Treatment abolishing LR bias** |
| --- | --- | --- | --- |
| (Pullar et al., 2001). | Human keratinocytes | 100mV/mm | 50nM KT 5720, 1μM ML-7, 15μM Sphingosine, 30μM W-7 |
| (Feng et al., 2012) | human neural stem cells (from H9 line) | 300mV/mm | 10μM Y27362, 5μg/mL AMD 3100, 25μg/mL AMD 3100 |
| (Yao et al., 2008) | rat hippocampal neurons | 300mV/mm | 100μM LY294002 |
| (Djamgoz et al., 2001) | Mat-LyLu | 3V/cm | 10μM veratridine |
|  | AT-2 | 3V/cm | 1μM TTX, 10μM veratridine |
| (Pu et al., 2007) | MTLn3 | 1V/cm | 2μM AG1478 |
| (Gao et al., 2011) | Dictyostelium | 12V/cm pH 6.5 | pH 5, pH 9 |

Interestingly, LR biases were also abolished by lowering the membrane potential of cultured cells via manipulation of the extracellular pH or ion concentration or altering the status of ion channels (Djamgoz et al., 2001; Gao et al., 2011). This finding is only predicted by the intracellular model, and is consistent with studies from a number of species including sea urchin, Xenopus, Ciona, zebrafish, and chick, which show that alterations in endogenous patterns of ion flux can disrupt embryonic LR patterning (reviewed in (Levin, 2006; Vandenberg and Levin, 2013)). It is notable that the functional data on asymmetry of migration indicates that single cells appear to utilize the same set of molecular components to establish consistent asymmetries as do whole embryos (cytoskeleton, motor proteins, ion channels) – a theme that has been echoed in other areas of cell biology (Marshall, 2011). The implication of ion flux in the asymmetry of human cell function further supports the idea that bioelectric signals have relevance to LR patterning beyond non-mammalian models (consistent with the conserved role of tubulin in asymmetry of plants, nematodes, and frogs (Lobikin et al., 2012)).

How might single cells utilize ion channels to establish consistent asymmetry? Recent developments in imaging of endogenous bioelectrical gradients have revealed that single cells do not have a single resting potential; rather, their plasma membranes contain multiple intracellular domains, where different regions of the cell are relatively hyper-or depolarized (Adams and Levin, 2012). It is conceivable that the interactions among these domains (and the resulting intracellular redistributions of charged molecules (Deloof, 1983; Jaffe, 1981; Jaffe et al., 1974; Lange and Steele, 1978; Larter and Ortoleva, 1981; Poo et al., 1979; Poo and Robinson, 1977; Poo et al., 1978)) establish within the cell an electrophoresis-like mechanism similar to what has been observed in the frog embryo (Adams et al., 2006; Esser et al., 2006; Fukumoto et al., 2005; Zhang and Levin, 2009). This “scale invariance” hypothesis for physiological mechanisms represents a fascinating area for future investigation, and suggests experiments that break drastically from the current paradigm focused on cilia hydrodynamics as the sole origin of vertebrate asymmetry.

### LR biases can be induced in otherwise unbiased cells

A small number of studies also revealed that cells that normally had unbiased responses to galvanotactic or chemotactic stimuli developed LR biases following additional manipulations (Table 3). For example, treatments that block ion transport proteins such as Ca^2+^ channels and Na^+^/K^+^-ATPases induced LR migration biases in keratinocytes, osteoblasts, neutrophils and osteosarcoma cells. Alterations inhibiting cytoskeletal remodeling also induced LR biases.

**Table 3.** Introduction of LR biases into unbiased cells

| Study | Cell type | Treatment lacking LR bias | Additional treatment inducing LR bias |
| --- | --- | --- | --- |
| (Trollinger et al., 2002) | Human Keratinocytes | 100mV/mm | 5mM Nickel |
| (Ozkucur et al., 2011) | fetal rat calvarial osteoblasts | 0.5 mV/mm | 10μM Oubain |
|  | SAOS-2 | 0.5V/mm | 5μM Oubain, 1μM PF573228 |
| (Allen et al., 1998) | Bac-1 Macrophages | CSF1 gradient | ~250μg/ml V12Cdc42 |
| (Malet-Engra et al., 2013) | Chronic lymphocytic leukemia cells | CCL19 | Knockdown of CIP4 (a Cdc42-interacting protein) |
| (Kumar et al., 2012) | mouse neutrophils WT | FMLP gradient | Wasp(-/-) [genetic mutant] |
| (Li et al., 2012) | MDCK I (monolayer) | 200mV/mm | 100nM PMA, 4mM EGTA in Calcium free medium, 4mM EGTA in normal medium, 50μg/mL DECCMA-1 |

We have previously described the presence of subtle asymmetries that are only revealed following functional perturbations including human genetic syndromes that unilaterally affect paired organs or structures (Pai et al., 2012). The developing somites, for example, appear symmetrical, but in fact have asymmetries that are ‘shielded’ by the action of retinoic acid signaling (Vermot and Pourquie, 2005). It is therefore plausible that in some cultured cells, ion transport proteins act to shield or compensate for asymmetrical gradients created by other pumps and channels. Thus, LR differences that are otherwise hidden can manifest when these ion transporters are inhibited.

The intracellular model of LR asymmetry predicts that asymmetry is a conserved feature of all cells. This theory is appealing because it is consistent with evolutionary theory, providing an intracellular origin of asymmetry that is conserved from the simplest of organisms up to humans. Our data identified chirality in a wide range of cells, implicating some of the same mechanisms used by embryos to orient the macroscopic LR axis. Consistent with recent data showing that individual cell chirality decisions control the large-scale asymmetry of multicellular structures grown in culture (Alpatov, 1946; Wan et al., 2011; Wan and Vunjak-Novakovic, 2011), it is reasonable to expect that evolution has leveraged the chiral properties of cells to drive asymmetric morphogenesis. Collectively, our analyses add new assays to complement recent findings that various forms of asymmetry occur in cultured cells and implicate avenues for future research that can link subcellular and cell-level asymmetries with the asymmetric organ placement observed in vertebrates and even the asymmetric behaviors (i.e. handedness) typical of humans.

## Methods

### Paper Inclusion Criteria

A literature search on galvanotaxis and chemotaxis was conducted using the databases PubMed and Scopus and separate search terms “galvanotaxis” and “chemotaxis”. Only analyses that included graphs showing either the migration trajectories or the final positions of at least 5 cells after they underwent galvanotactic or chemotactic migration were included in the final analyses.

### Defining the LR axis

The original graphs were most frequently wind rose plots; the direction of electric fields and chemical gradients were clearly labeled, and each cell’s initial position was superimposed onto a common origin point, the center of either a Cartesian or angular coordinate system (Fig. 1). The direction of cell movement toward the attractant defined one “axis”, and the source of the attractant was arbitrarily labeled “anterior”. The substrate surface was arbitrarily considered the “ventral” side of the cells. The axis pointing toward the target of cell migration, such as a source of chemoattractant or the electrode to which cells of a certain type tend to travel, was used as the midline about which left and right were defined (Fig. 1). Only treatments that induced a positive response (i.e. movement toward the electric or chemical attractant) were used in the LR analysis.

### Quantifying Leftward and Rightward Migration

Only cells considered sensitive to their electric field or chemical gradient (i.e. if their final positions were in the half of the graph closest to the attractant) were assessed for a LR bias. In graphs in which each cells’ final position was represented by a mark, the number of cells travelling right or left was determined by counting the number of marks in the quadrant lying to the right or left of the midline, respectively.

In graphs in which individual cells’ migration trajectories were shown, it was often difficult to determine the final positions of individual cells. In this circumstance, GIMP 2 photo processing software was used to count the number of pixels lying to the left and right of the axis defining the midline. GIMP was also used to analyze graphs in which the marks of cells’ final positions were too close together to count definitively.

### Statistical Analysis of LR Bias

A Chi-Square test was conducted to test the observed numbers of left and right cells/pixels against a null hypothesis in which 50% of cells/pixels lay to the left of the midline and 50% lay to the right. Data from multiple graphs within a single paper were condensed if cells were of the same type and treated identically including electric field strength, type of chemoattractant, drug dosage, etc. Cases were considered to show significant LR bias when p <0.05.

## Author Contributions

KGS identified the studies to be incorporated in the final analyses, collected the data from individual studies and built the original data tables. KGS, LNV and ML analyzed the data and wrote the manuscript. LNV created the figure and tables.

## Acknowledgements

The authors thank Dr. Daniel Lobo for assistance in the use of GIMP software program, and members of the Levin Lab for helpful discussions on these findings. This work was funded by the American Heart Association Established Investigator grant 0740088N and NIH grants R01-GM077425 (to ML) and NRSA grant 1F32GM087107 (to LNV). M.L. is also supported by the G. Harold and Leila Y. Mathers Charitable Foundation.

